# DNA crookedness regulates DNA mechanical properties at short length scales

**DOI:** 10.1101/283648

**Authors:** Alberto Marín-González, J. G. Vilhena, Fernando Moreno-Herrero, Rubén Pérez

## Abstract

Sequence-dependent DNA conformation and flexibility play a fundamental role in specificity of DNA-protein interactions. Here we quantify the DNA crookedness: a sequence-dependent deformation of DNA that consists on periodic bends of the base pair centers chain. Using molecular dynamics simulations, we found that DNA crookedness and its associated flexibility are bijective: unveiling a one-to-one relation between DNA structure and dynamics. This allowed us to build a predictive model to compute DNA stretching stiffness from solely its structure. Sequences with very little crookedness show extremely high stiffness and have been previously shown to form unstable nucleosomes and promote gene expression. Interestingly, the crookedness can be tailored by epigenetic modifications, known to affect gene expression. Our results rationalize the idea that the DNA sequence is not only a chemical code, but also a physical one that allows to finely regulate its mechanical properties and, possibly, its 3D arrangement inside the cell.

## MAIN TEXT

The mechanism by which proteins interact with the genome with such extraordinary specificity is still an open question in biology. Since the discovery of the DNA double helix (dsDNA), it became clear the existence of a sequence dependent set of hydrogen bond donors and acceptors that are exposed in the major groove and specifically recognized by certain amino acids. However, there is increasing evidence that this mechanism is far from sufficient. In a number of DNA-protein complexes, DNA adopts a conformation that substantially deviates from the canonical B-form (*1-3*), suggesting a structural deformation or an exceptional flexibility intrinsic to the DNA sequence. Among the most-studied cases are sequence-dependent DNA deformations –A-like structures, kinked base pair steps and A-tracts– that play an important role in transcription regulation (*4-7*). In parallel, the high sequence-dependent flexibility of DNA is used by several proteins to achieve binding specificity (*1, 8*).

However, many aspects of DNA flexibility have so far remained elusive. Indeed, It is not fully understood how a relatively stiff molecule, with a persistence length of P∼50 nm, is able to wrap around a histone octamer of ∼4 nm of radius. Even more intriguing is the fact that some sequences are hardly able to form stable nucleosomes, arguably as a consequence of a distinct conformation or mechanical properties (*9, 10*). The same question holds for other DNA-protein complexes, in particular, for many repressor systems where a highly bent loop is predicted in the DNA (*11*). These considerations, supported by recent findings on high bendability of DNA at short-length scales (*12, 13*) challenge the currently accepted worm-like chain model (WLC) and demand for an accurate description of sequence-dependent DNA flexibility at these scales.

Using constant-force molecular dynamics (MD) simulations (*14*) we observed that the extension of the DNA changed from one sequence to another for molecules with the same number of base pairs. we observed that the extension of the DNA changed from one sequence to another for molecules with the same number of base pairs. We performed over 1 μs-long MD simulations of 18 base pair long DNA molecules with benchmark sequences of the form CGCG(NN)_5_CGCG, where NN denotes AA, AC, AG, AT, CG and GG, and computed its average structure at 1 pN force, see **Fig. 1A** and **fig. S1**. The variability in the extension could not be attributed to a different separation between consecutive base pairs (**fig. S2**) and, therefore, reflects an intrinsic curvature of the molecule. This curvature is apparent if we represent the centers of the base pairs (color beads in **Fig. 1A** and **fig. S1**). We will denote this curvature by crookedness, in analogy with a crooked road, whose trajectory is not straight. Importantly, DNA crookedness is of the order of a few (∼ 2) nanometers, a scale comparable with the histone octamer radius (*15*), the DNA curvature of the DNA-I-PpoI endonuclease complex (*16*) and several examples of sharply bent DNA found in regulatory regions (*11*). **Figure 1B** illustrates this point, comparing the large ∼16nm radius of curvature predicted by the WLC model for a 30 bps DNA molecule with the crookedness, that provides the local curvature of few nm, necessary to bend over a nucleosome core or wrap around a protein complex.

**Figure 1.**
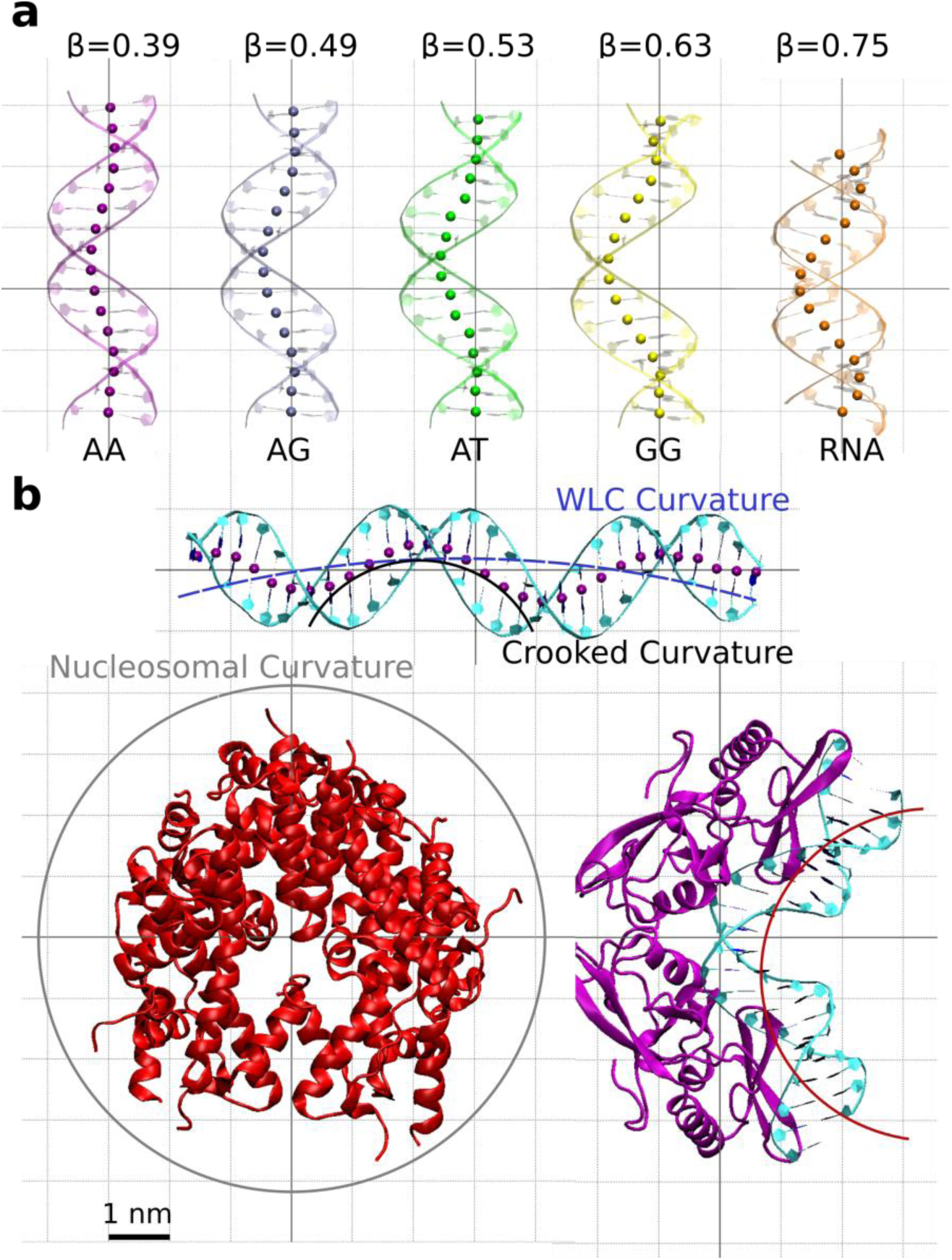
Representation of DNA crookedness. **a**, Average structures and computed β values (in rads) over 1 μs-long MD simulation at 1 pN force of the sequences CGCG(NN)_5_CGCG with NN=AA, AC, AT and GG. The beads represent the centres of the base pairs. The terminal base pairs have been omitted. **b**, (top), average structure over 250 ns MD of a 30 bps poly-G DNA molecule. The solid black line represents the crookedness curvature and the dashed blue line an estimation of the curvature predicted by the WLC. (bottom), examples of highly curved DNA when bound to proteins. (left), histone octamer crystallized in (*15*) (PDB ID: 1AOI), where the histone tails have been removed for clarity. A grey circle of radius 41.8 Å represents the trajectory of nucleosomal DNA. (right), crystal structure of the homing endonuclease I-PpoI DNA complex taken from (*16*) (PDB ID: 1A73), with an estimation of the DNA curvature represented by the solid red line.

Crookedness can be quantified via a parameter *β* defined as cos *β* ≡ *x* / Σ*l*_*i*_, being *x* the extension (end-to-end distance) of the molecule and Σ*l*_*i*_ the sum of distances between consecutive base pair centers. When the molecule is completely straight the line that runs through the base pairs is perfectly aligned, *x* = Σ*l*_*i*_, and the crookedness, β, is zero. As this line deviates more from the helical axis, the ratio *x* / Σ*l*_*i*_, becomes smaller and therefore DNA crookedness increases. Double stranded RNA (dsRNA), which is A-form, is the most crooked, whereas B-DNA molecules exhibit lower β values centered on 0.5 (**Fig. 1A.** Indeed, as shown in **fig. S3**, the crookedness is a reasonable parameter to distinguish DNA conformations along the A ↔ B spectrum.

Additionally to sequence-dependent conformations, proteins often exploit DNA flexibility (*1, 8*). Therefore, a complete comprehension of the biological relevance of DNA crookedness requires understanding its effect on DNA mechanical properties. We propose a model to rationalize the relation between β and DNA stretch modulus, *S*, where DNA can elongate by reducing its β or by separating consecutive base pairs (**Fig. 2A, B** and supplementary material). Thus can be written for an N-bps molecule as a set of N springs in

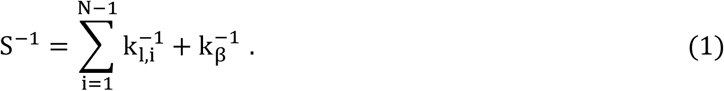

**Figure 2.**
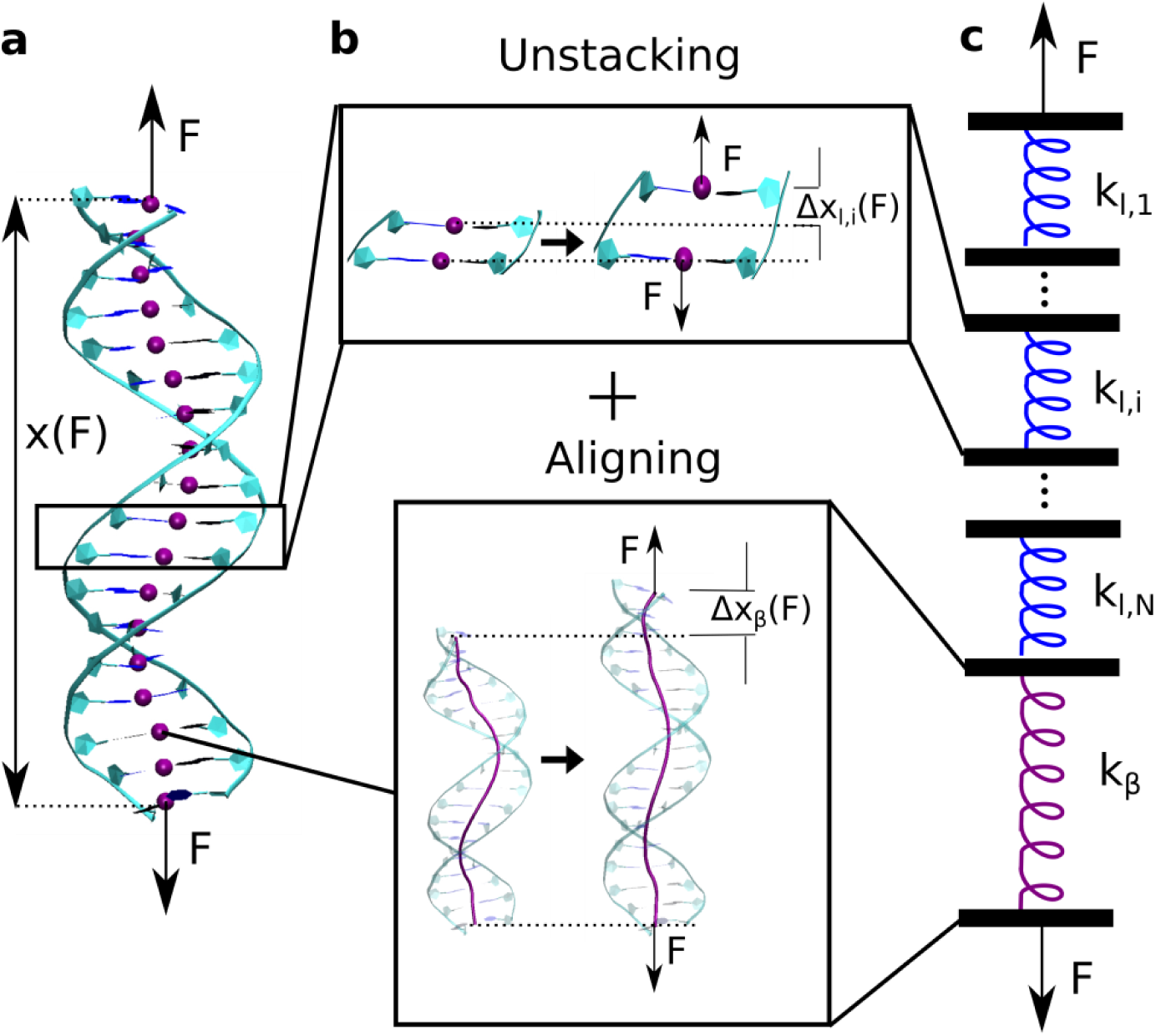
Model to relate DNA crookedness with the stretch modulus, *S*. **a**, DNA molecule with the base pair centers represented by purple beads. An external force, *F*, induces a change in DNA extension, *x*(*F*) **b**, DNA can elongate by separating consecutive base pairs (top; Δ*x*_*l,i*_) or by aligning the base pair centers with the helical axis (bottom; Δ*x*_*β*_). **c**, DNA is modelled as a set of N springs in series. The first N-1 springs account for the stiffness of elongating individual base pair steps,, and the N^th^ spring is the crookedness stiffness, *k*_*β*_.

The first N-1 springs correspond to the stiffness of separating consecutive base pairs, denoted by *k*_*l,i*_, and the N^th^ spring to the crookedness stiffness, denoted by *k* _*β*_ (**Fig. 2C** and **fig. S4**).

We ran five constant-force MD simulations at F = 1, 5, 10, 15 and 20 pN using the benchmark sequences described above to determine the sequence dependence of *k*_*l,i*_ and *k*_*β*_. We found that consecutive base pair elongation (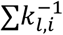, **Fig. 2C** blue springs) has a minor contribution to (f**ig. S5, S6**) and the dominant contribution comes from *k*_*β*_, which accounts for a *global* deformation of the molecule as a whole (**Fig. 2C** purple spring). We computed *k*_*β*_ by measuring the force induced change in DNA crookedness, Δcos*β(F)*, according to (supplementary material):

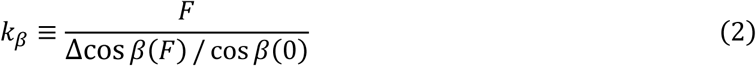

**Figure 3A** shows the calculated *k*_*β*_ values (blue points) and the extraordinary fit provided by the function *k*_*β*_(*β*) = Ae^-*kβ*^ + *B* with parameters and *A* = (2.24 ± 1.24) ×10^6^ *pN* and *k* = 16.2 ± 1.5. Remarkably, this function reproduces the results for 11 additional sequences exhibiting a broad range β of values (table S2). This reveals an exceptional property of DNA crookedness: the relation with its associated stiffness is bijective. In other words, an *equilibrium* structural parameter, β, univocally determines a *dynamical* response, *k* _*β*_; and vice versa.

**Figure 3.**
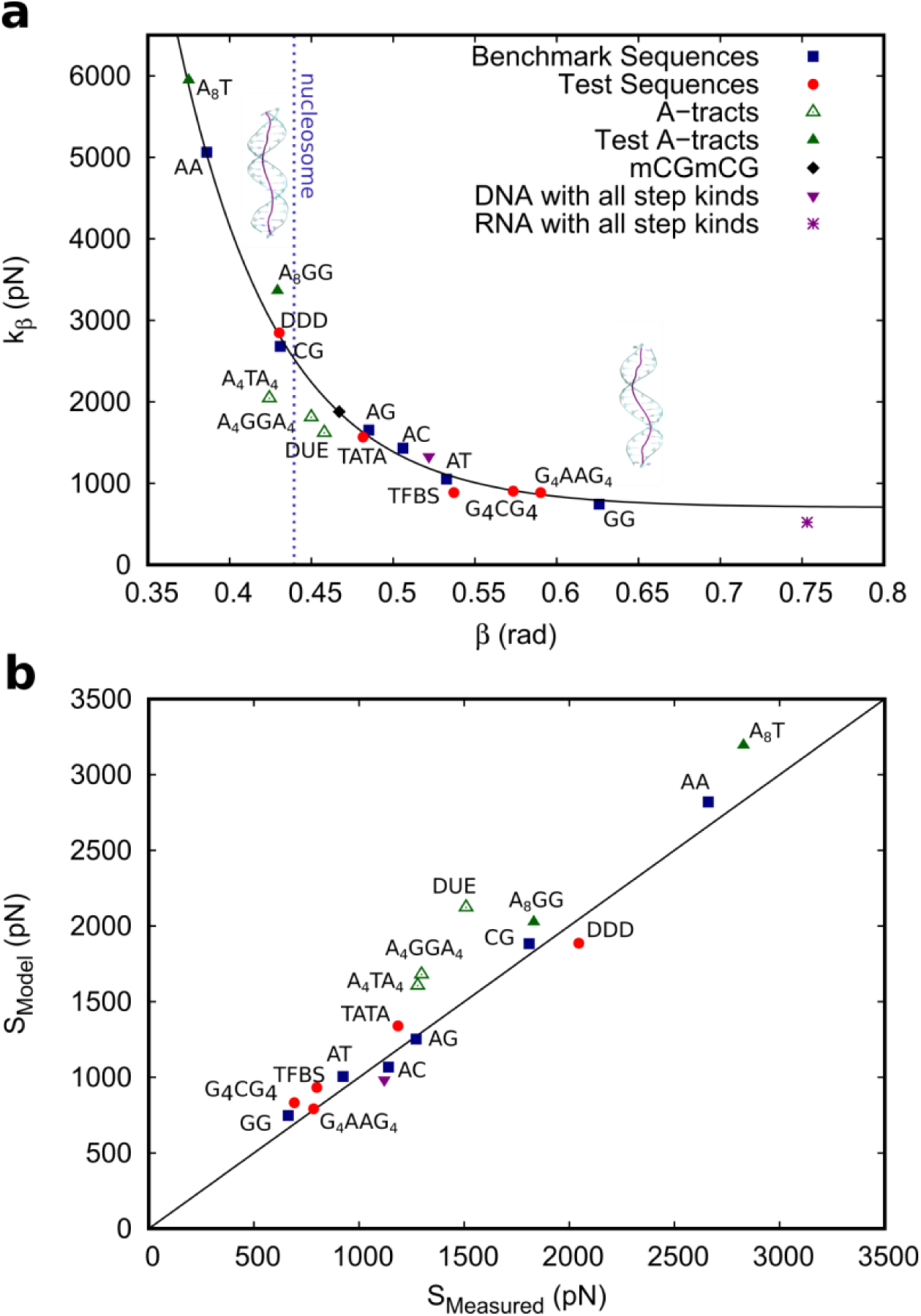
Implications of DNA crookedness on DNA flexibility. **a**, *k*_*β*_ values as a function of the crookedness, *β*. *k*_*β*_ was computed from the MD simulations data using **Eq. 2** and taking the F = 1 pN simulation as reference. *β* was computed from the F = 1 pN simulation as 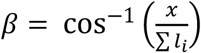. The blue squares correspond to the benchmark sequences and are the points used for the fit *k*_*β*_(*β*) = Ae^-*kβ*^ + *B* (fitting parameters in main text). The red circles represent the sequences simulated to test the model. The void green triangles are the A-tracts containing sequences and the filled green triangles are the sequences used to test the effect of the A-tracts. See **table S3** for a list of the simulated sequences. The dashed line represents an estimation of nucleosome curvature according to (*28*). **b**, The computed value of is plotted as a function of the value directly measured from the force-extension curve as done in (*14*). The stretch modulus was computed from our model for the simulated sequences described above using **Eq. 1**, the value of the fit of *k*_*β*_, **table S1** and the values of *β* and Σ*l*_*i*_ obtained from the F = 1 pN simulation.

The extraordinary one-to-one correspondence between *β* and *k*_*β*_ provides predictive power to our model. Indeed, if the equilibrium structure of a DNA sequence is known (by NMR, crystallography, MD…) one can measure *β* and the base pair separation distances. Then, *k*_*β*_ can be obtained from **Fig. 3A** and the *k*_*l,i*_ taken from **table S1**, to compute the value of *S* with **Eq. 1. Figure 3B** confirms the perfect agreement with the measured values –our calculations and refs (*17, 18*)-for a wide range of values (∼800-3000 pN).

In addition to predicting the stretch modulus, our model provides valuable information about the nature of its sequence dependence. Our results show that *S* is not dependent on the GC content, but rather, on how guanines and cytosines are distributed along the strands. Molecules with alternating GC’s are much less flexible than those where several guanines are placed sequentially on the same strand. This confirms the idea that mechanical stability is independent of thermal stability, as suggested in experiments of plectoneme formation (*19*). Moreover, our model identified three molecules with anomalously enhanced flexibility (**Fig. 3A** void green triangles). These molecules include two strings of three or four A-tracts, which are known to introduce anomalies in DNA curvature and mechanical properties (*20, 21*). The possibility that this enhanced flexibility stems from the presence of two A-tracts in A_4_TA_4_ and A_4_GGA_4_ molecules was confirmed by running additional simulations using the sequences A_8_T and A_8_GG (**Fig. 3A** filled green triangles). This highlights that other sources of flexibility may coexist with the crookedness mechanism proposed here.

Our benchmark simulations allowed us to identify sequence patterns with unusually high stiffness, namely poly-A and alternating CG. They appear repeatedly throughout the genome and are frequently involved in gene expression regulation (*9, 10*). Long poly(dA:dT) and their flanking DNA have been shown to be depleted of nucleosomes *in-vitro* suggesting a mechanism of gene activation (*9*). In parallel, about 70% of annotated gene promoters are associated with so-called CpG islands (CGI), rich in CpG steps (*10*). Notice that our CG-alternating sequence shows the largest proportion of CpG steps possible in any molecule: one in every two steps. As in the case of the poly-A, CGI have been attributed to nucleosome destabilization (*22*). These findings together with **Fig. 3A** suggest a possible relation between an unusually high crookedness stiffness and nucleosome destabilization. Hypermethylation of CGI commonly induces gene silencing and in some cases this has been attributed to nucleosome stabilization (*10*). We found that complete hypermethylation of the poly-CG molecule significantly increases its crookedness and flexibility, in quantitative agreement (table S3) with recent optical tweezers experiments reporting both an unusually high *S* and a softening induced by hypermethylation in CGI (*23*). This could increase nucleosome affinity for a hypermethylated CGI (*24*). Notice however, that outside this context, methylation is known to reduce DNA flexibility and destabilize nucleosomes (*25-27*).

We have shown that DNA crookedness can be tuned by specific sequences, being the main responsible for a sequence dependent stretching flexibility. It is tempting to extend this feature to explain the high DNA bendability reported at short scales (80-100 bps) (*12, 13*). DNA could bend around the histone octamer by periodically stretching segments at the length scale of DNA crookedness (**Fig. 4A**). The resulting overall DNA elongation would be in agreement with the finding that nucleosomal DNA is stretched by 1-2 bps (*28*).

**Figure 4.**
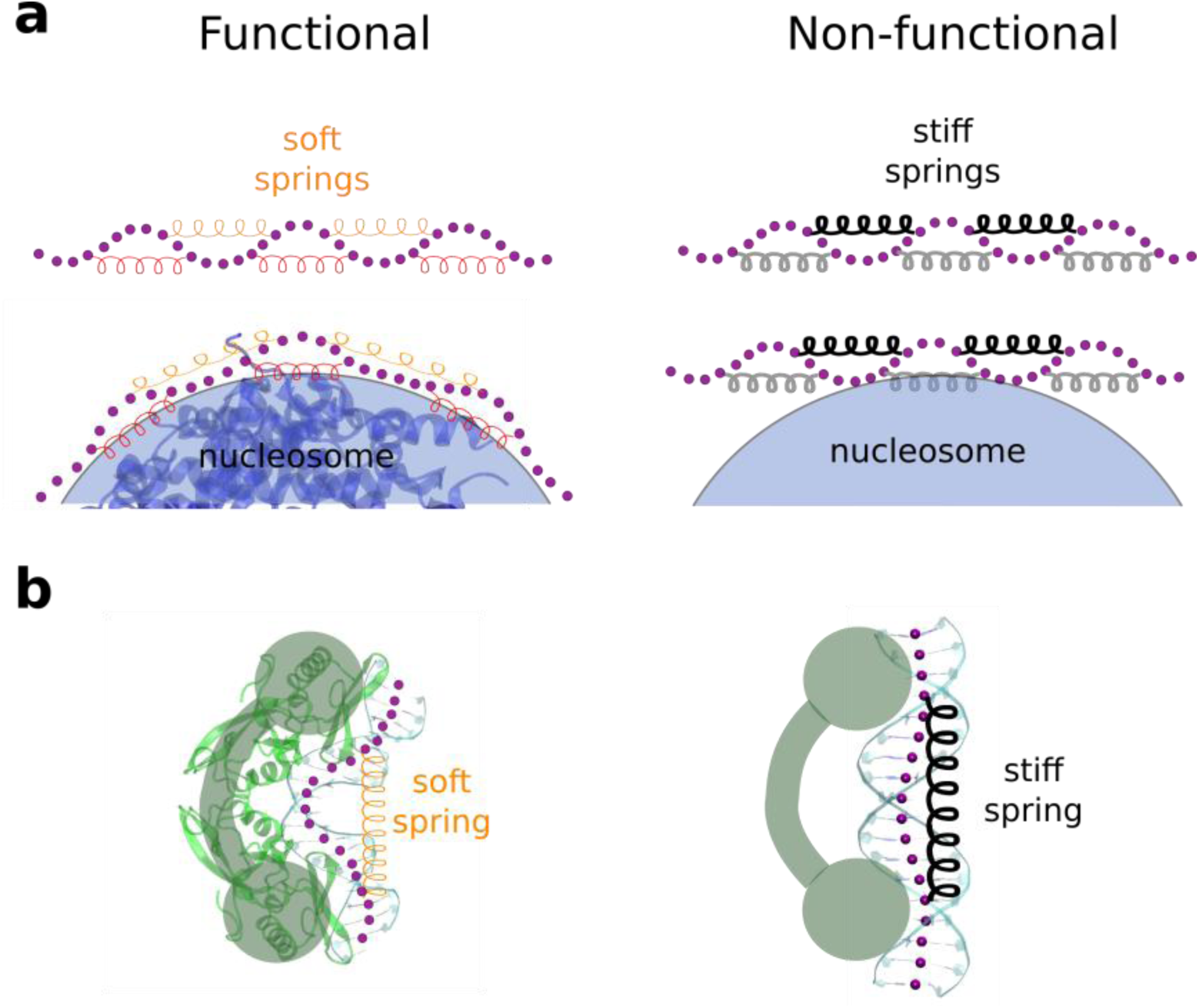
Implications of DNA crookedness on DNA-protein interactions. **a**, The base pair centres chain of a DNA molecule is represented by purple beads and the crookedness flexibility by springs. For flexible molecules (left) short-scale bending can be achieved by elongating the upper springs. This mechanism would be precluded for molecules with low flexibility (right) **b**, crookedness flexibility could also be exploited to induce a conformational change in DNA towards an A-like structure. On the left is the crystal structure of the I-PpoI DNA complex (*16*) (PDB ID: 1A73), the same as in **Fig. 1B**. When bound to this protein a high distortion is found on the DNA, which would be more favourable for highly crooked and flexible sequences (left) than for sequences for which this flexibility is hindered (right).

Importantly, this crookedness regulation would be energetically more costly for molecules that are exceptionally stiff, such as the poly-A, in line with the observation that long (>20 bp’s) A-tracts present a slower looping rate than a random sequence at length scales below the persistence length (*13*). This seems to us a reasonable mechanism of transcription regulation based on DNA crookedness. On the other hand, highly crooked molecules might be preferred for DNA-ligand binding where DNA flexibility is required. Indeed, interaction of DNA with proteins and drugs commonly modifies the B-DNA to a more A-like form and this occurs in a sequence-dependent manner (*3*). According to **Figure 2** molecules with high *β* would be more prone to form an A-type helix when bound to a protein both because their structure is already closer to the A-form and because of their enhanced flexibility (**Fig. 4B**). Additionally, crookedness may modulate the charge distribution along the duplex, known to be a mechanism for protein-DNA recognition by electrostatic interactions (*7*). Altogether, our results provide evidence for the presence of a ’*hidden code*’ imprinted in the genome that modulates its 3D structure and might contribute to gene expression regulation. The extent to which this plays an active role, allowing structural deformations of the naked DNA; or a passive one, where the proteins exploit this flexibility to induce non-canonical DNA conformations will be a subject of future studies.

## ACKNOWLEDGMENTS

We thank the financial support from the Spanish MINECO (projects MAT2014-54484-P, FIS2014-58328-P, MAT2017-83273-R (AEI/FEDER, UE), and BFU2017-83794-P (AEI/FEDER, UE)). F.M.-H. acknowledges support from European Research Council (ERC) under the European Union’s Horizon 2020 research and innovation (grant agreement No 681299). A.M.-G. acknowledges support from the International PhD Program of “La Caixa-Severo Ochoa” as a recipient of a PhD fellowship. The authors acknowledge the computer resources, technical expertise and assistance provided by the Red Española de Supercomputación at the Minotauro Supercomputer (BSC, Barcelona). We thank J. Lipfert, G.J.L. Wuite, C.L. Pastrana and A. Gil for fruitful discussions.

## AUTHORS CONTRIBUTIONS

A.M.-G. and J.G.V. performed MD simulations and data analysis. R. P. and F. M.-H. designed simulations and supervised research. All authors contributed to the writing and revision of the paper.

## ADDITIONAL INFORMATION

This article contains Supporting Information.

